# Distinct Neural Representations of Decision Uncertainty in Metacognition and Mentalizing

**DOI:** 10.1101/2021.05.27.445947

**Authors:** Shaohan Jiang, Sidong Wang, Xiaohong Wan

**Affiliations:** State Key Laboratory of Cognitive Neuroscience and Learning and IDG/McGovern Institute for Brain Research, Beijing Normal University, Beijing, 100875, China

**Author notes:** Corresponding author **Email:** (XW). These authors contributed equally to this work (Shaohan Jiang, Sidong Wang). **Author Contributions:** S. Jiang and S. Wang conducted the experiments; S. Jiang and X. Wan analyzed the data; S. Wang and X. Wan designed the experiments; X. Wan wrote the manuscript and supervised the project.

**Keywords:** mentalizing, metacognition, theory of mind, decision uncertainty, dorsomedial PFC

## Abstract

Metacognition and mentalizing are both associated with meta-level mental state representations. Specifically, metacognition refers to monitoring one’s own cognitive processes, while mentalizing refers to monitoring others’ cognitive processes. However, this self-other dichotomy is insufficient to delineate the two high-level mental processes. We here used functional magnetic resonance imaging (fMRI) to systematically investigate the neural representations of different levels of decision uncertainty in monitoring different targets (the current self, the past self, and others) performing a perceptual decision-making task. Our results reveal diverse formats of intrinsic mental state representations of decision uncertainty in mentalizing, separate from the associations with external information. External information was commonly represented in the right inferior parietal lobe (IPL) across the mentalizing tasks. However, the meta-level mental states of decision uncertainty attributed to others were uniquely represented in the dorsomedial prefrontal cortex (dmPFC), rather than the temporoparietal junction (TPJ) that also equivalently represented the object-level mental states of decision inaccuracy attributed to others. Further, the object-level and meta-level mental states of decision uncertainty, when attributed to the past self, were represented in the precuneus and the lateral frontopolar cortex (lFPC), respectively. In contrast, the dorsal anterior cingulate cortex (dACC) consistently represented both decision uncertainty in metacognition and estimate uncertainty during monitoring the different mentalizing processes, but not the inferred decision uncertainty in mentalizing. Hence, our findings identify neural signatures to clearly delineate metacognition and mentalizing and further imply distinct neural computations on the mental states of decision uncertainty during metacognition and mentalizing.

## Introduction

Humans are social beings. We interact with others not only in the physical world but also in the mental world. Differing from objects, humans are free and intentional agents who can hold their mental states that are not direct reflections of reality. The human brain thus needs to concurrently represent mental states reflecting the physical world and the mental worlds of both the self and others [1–3]. Failures of normal development of such an ability can cause deficits in human cognition and behaviors, for instance in autism spectrum disorder (ASD) and schizophrenia [4, 5]. Thus, it is a central question in psychology and neuroscience to understand the mechanisms of human mental state representations.

A principal criterion to distinguish non-social activities from social activities is whether the activity is conducted towards the self or others [6–9]. A corresponding distinction is also drawn for mental state attributing processes: monitoring one’s own cognitive processes is referred to as metacognition [10], but when the target subject is an intentional agent other than the self it is referred to as mentalizing [11–13]. Although both metacognition and mentalizing involve meta-representations of the mental world [1–3], the representational formats and sources of the mental states differ. Critically, mentalizing necessitates others’ perspective taking to infer their mental states [14], while one’s own mental states are directly accessible in metacognition [1, 2]. On the other hand, the mental states attributed to others might be also inferred through associations between external information and internal mental states. Thereby, it is difficult to discern the underlying processes merely from the observed behaviors [15–17]. Because of this ambiguity, to date, it remains unclear whether or not non-human primates can mentalize [15, 17].

However, the self-other dichotomy on the target agents is insufficient to fully discern the two processes. For instance, similar to attributing mental states to others, the mental states of the past self are also inaccessible in that the available information is only from external cues. That is, the momentary mental states of the past self cannot be inspected as in metacognition but can be only inferred. It thus becomes ambiguous whether the mental state representations are similar to metacognition or mentalizing. Like the two-order hierarchy of type 1 (object-level) and type 2 (meta-level) mental states and processes in metacognition [10], the mental states attributed to others in mentalizing could also be hierarchically categorized (Fig 1). Thereby, the representations of the two-level mental states even for the same target in mentalizing might be separate. Further, during monitoring others’ object-level performance, the evoked meta-level mental states are actually tied to the observer, rather than the target subjects. In this sense, the mental state representations are more similar to those in metacognition than mentalizing. Hence, in many situations the distinctions between metacognition and mentalizing along the self-other dichotomy are ambiguous. Therefore, we looked to neural signatures to clearly delineate metacognition and mentalizing.

**Fig 1.**
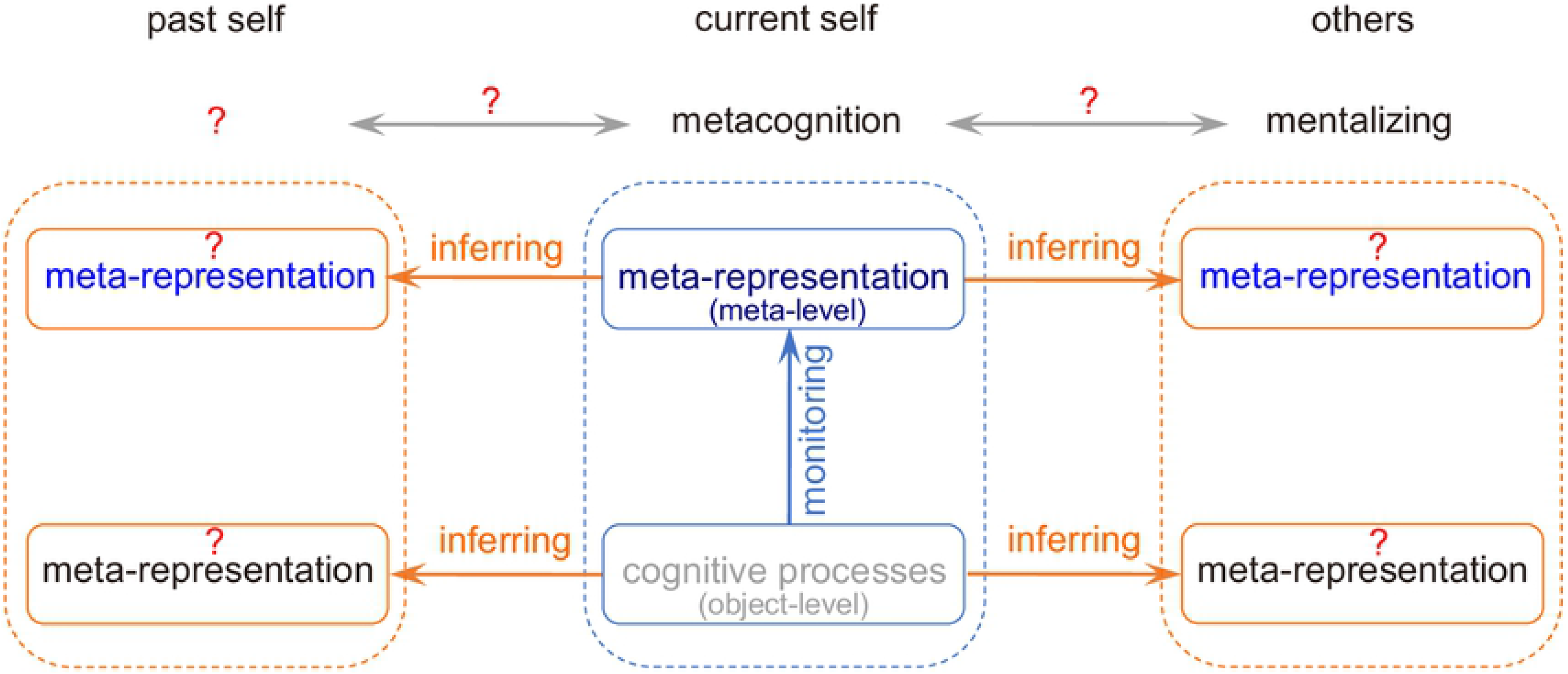
The schematic structure of meta-representations in metacognition and mentalizing. Within the cognitive control framework, the meta-representations in metacognition are internally-generated meta-level mental states generated through introspecting or monitoring the object-level cognitive processes performed by the current self (center panel). When mentalizing about the other, the meta-representations of the object-level and meta-level cognitive processes attributed to the other are inferred because the other’s internal processes are inaccessible (right panel). However, the self-other dichotomy between metacognition and mentalizing cannot determine whether the process of attributing mental states to the past self is metacognition or mentalizing (left panel). The current study is aimed to investigate these meta-representations in the human brain and the relationship between metacognition and mentalizing.

Surprisingly, although a number of disparate studies on the neural mechanisms of metacognition and mentalizing have been conducted in cognitive neuroscience [18, 19] and social neuroscience [20, 21], respectively, a direct comparison of the two neural processes is so far lacking. This situation might be primarily due to the lack of an appropriate experimental paradigm applicable for both processes. The mental state that is mainly addressed in studies of metacognition is decision uncertainty (the meta-level mental state), that is, the extent to which one subjectively believes that one’s own decision is incorrect (decision inaccuracy, the object-level mental state). Decision uncertainty is an endogenous control signal for improving one’s decision even with no external feedback [18, 22]. Importantly, it also serves as a critical social signal for efficacious decision improvement in joint decision-making [23, 24]. Hence, it is of great importance to understand how to attribute the mental states of decision uncertainty to the target subjects other than the current self in mentalizing.

In the current study, we aimed to delineate the neural representations of different levels of decision uncertainty attributed to different target subjects: the current self, the past self and others. To reveal the relationship between metacognition and mentalizing, we adapted a task paradigm often used in metacognition to apply to mentalizing. We used functional magnetic resonance imaging (fMRI) to characterize the neural representations of object-level (type-1) and meta-level (type-2) intrinsic mental states of decision uncertainty attributed to others and the past self in mentalizing, separate from the associations with external information. We found diverse mental state representations in the mentalizing tasks. First, external information was commonly represented in the right inferior parietal lobe (IPL) across the mentalizing tasks. Second, the mental states of decision uncertainty attributed to others were uniquely represented in the dorsomedial prefrontal cortex (dmPFC), but not in the temporoparietal junction (TPJ) that also equivalently represented the decision inaccuracy attributed to others. Third, the type-1 and type-2 mental states of decision uncertainty, when attributed to the past self, were represented in the precuneus and the lateral frontopolar cortex (lFPC), both also involved in metacognition. In contrast, the dorsal anterior cingulate cortex (dACC) selectively represented decision uncertainty in metacognition and estimate uncertainty during monitoring the mentalizing processes, but not the inferred decision uncertainty in mentalizing. Therefore, our findings demonstrate distinct formats of mental state representations in mentalizing but a general format of mental state representations in metacognition.

## Results

### Task paradigm

We carried out three fMRI experiments investigating the mental state representations of decision uncertainty in metacognition and mentalizing (Fig 2a). Twenty-eight healthy participants took part in all of the experiments (see Methods). In experiment 1, the participant judged the gross motion direction of random moving dots and rated her subjective uncertainty about the decision (Fig 2b). There were four different task difficulty levels randomly mixed in the task (S1 Fig). Hereafter this perceptual decision-making task was referred to as the random dot motion (RDM) task (Fig 2a). The participant reported the current-self mental states of decision uncertainty (CS-DU) via metacognition immediately accompanying perceptual decision-making in each trial, that is, “ *I am this uncertain in **my** decision*.”

**Fig 2.**
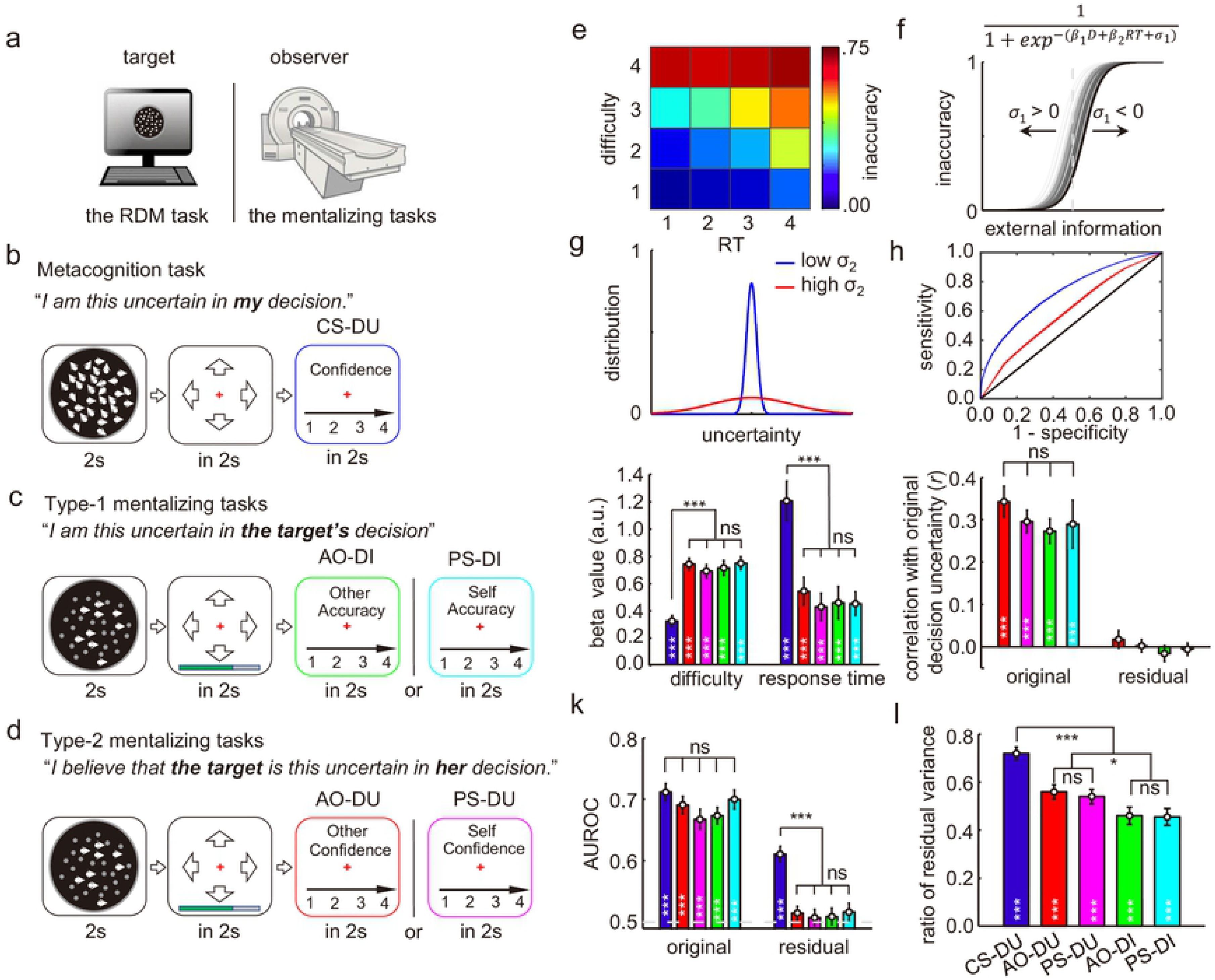
Task paradigms and behavioral results. (**a**) Task setup for the mentalizing tasks. The participant inside the MRI scanner observed the random dot motion (RDM) task performance done concurrently by the target subject who was outside the scanner. (**b**) The metacognition task: The participant completed the RDM task and reported her current-self decision uncertainty (CS-DU). (**c**) The type-1 mentalizing tasks: The participant observed the RDM task performance by a target subject and reported the target subject’s decision inaccuracy. The target was either an anonymous other (AO-DI) or the past self (PS-DI). (**d**) The type-2 mentalizing task: Instead of judging the target subject’s decision inaccuracy, the participant estimated the decision uncertainty that would be concurrently reported by the AO/PS in the current trial (AO-DU/PS-DU). (**e**) The decision inaccuracy empirically changed with the task difficulty and reaction time (RT) in the AO-DU task, averaged across participants. (**f**) Decision inaccuracy is a sigmoid function of task difficulty and RT on each trial. Each target subject has unique internal noise (*σ*_1_) in perceptual decision-making that causes variances in decision inaccuracy. (**g**) In estimating the target subject’s decision uncertainty, the unique internal noise (*σ*_2_) needs to be considered in mapping decision inaccuracy to decision uncertainty. (**h**) Different levels of internal noise (*σ*_2_) cause different metacognitive abilities (area under the ROC curve, AUROC). (**i**) The regression beta values of the normalized (task) difficulty and RT in each task. They did not differ across mentalizing tasks (ANOVA, task difficulty: F_[3,112]_ = 0.11, *P* = 0.95; RT: *F*_[3,112]_ = 0.16, *P* = 0.92), but significantly differ from the metacognition task (two-tailed paired t-test, task difficulty: t_27_ = 6.2; *P* = 2.5 × 10^−8^; RT: t_27_ = 4.1; *P* = 5.9 × 10^−5^). (**j**) The correlation between the estimated decision inaccuracy/uncertainty in the mentalizing tasks with the target’s original decision uncertainty reported in the metacognition task, before (original: ANOVA, F_[3,112]_ = 0.28, *P* = 0.84) and after (residual: *P*s > 0.30) the associations with external information were regressed out. (**k**) The prediction consistency between the decision outcome (true or false) and the reported/estimated decision inaccuracy/uncertainty measured by the AUROC, before (original: ANOVA, F_[4,139]_ = 1.22, *P* = 0.31) and after (residual: *P*s > 0.20) the associations with the external information were regressed out. (**l**) The ratio of the estimated residuals to the total estimated variances after the associations with external information were regressed out (two-tailed t-test, AO: t_27_ = 2.5; *P* = 0.0096 in the contrast between type-2 and type-1 mentalizing tasks; PS: t_27_ = 2.1; *P* = 0.023 in the contrast between type-2 and type-1 mentalizing tasks; two-tailed paired t-test, t_27_ = 3.5; *P* = 3.3 × 10^−4^ in the contrast between the metacognition task and the mentalizing tasks). The error bars represent standard error of the mean (SEM) across participants. \**P* < 0.05; \*\**P* < 0.01; \*\*\**P*< 0.001.

In experiment 2, the participant inside the scanner observed an anonymous other (AO) concurrently performing the RDM task outside the scanner and judged the AO’s decision inaccuracy (AO-DI). That is, “*I am this uncertain in **the target’s** decision*” (Fig 2c). Differing from the metacognition task (CS-DU), the AO’s cognitive states were inaccessible and necessitated the participant to infer. To avoid evoking the participant’s own decision uncertainty, the stimuli presented to the participant were noiseless: Only coherently-moving dots were moving, whereas randomly-moving dots remained stationary. By virtue of this altered stimulus presentation, the participant could perceive the task difficulty without evoking her own decision uncertainty (Fig 2c). This was a necessary condition to dissociate the neural representations of decision uncertainty in mentalizing from those in metacognition. Further, the AO’s response time (RT) was reported to the participant by a progress color bar, whereas neither the AO’s choice nor the reported decision uncertainty was presented to the participant. Hence, the participant could only use the external information of the task difficulty and the RT to estimate the AO’s decision inaccuracy. In a parallel task, the participant alternatively observed performance on the RDM task previously done by the self and judged the past-self decision inaccuracy (PS-DI). Otherwise, the experimental procedure was identical to the AO-DI task. Notably, as the decision variables of the past self were also inaccessible, the underlying process in both tasks should be mentalizing. We refer to these tasks as the type-1 mentalizing tasks.

In experiment 3 (Fig 2d), the experimental procedure was identical to the type-1 mentalizing tasks (experiment 2), but the participant estimated the AO/PS’s meta-level mental states of decision uncertainty in each trial (AO-DU and PS-DU). That is, “*I believe that **the target** is this uncertain in **her** decision*.” The two tasks thus also entailed mentalizing to attribute mental states of decision uncertainty to the AO/PS. We refer to these two tasks as the type-2 mentalizing tasks.

The task sequences of all the mentalizing tasks were identical, only the instructions differed. Thereby, any behavioral and neural differences between them should be caused by different mentalizing processes. The task sequences of the metacognition task and the mentalizing tasks were also quite similar. However, the differences between the two types of tasks existed in both the perception (object-level) phase and judgment (meta-level) phase. We mainly concerned here with the latter phase. The participant monitored internally-generated decision uncertainty in the metacognition task and inferred decision uncertainty in the mentalizing tasks. Our simulation analyses demonstrated that the neural correlates of the reported decision uncertainty in either the stimulation phase or the judgment phase were recoverable by the conventional general linear models (GLMs) with the parametric modulation effects in both phases (S2 Fig). Further, we also segregated the intrinsic mental states of decision uncertainty from the components directly associated with perception (task difficulty and RT) in both types of tasks.

### Hierarchical representations of decision inaccuracy and decision uncertainty

According to the decision-making theories [25], decision inaccuracy is crucially dependent on both task difficulty and RT (Fig 2e). The higher the task difficulty and the longer the RT, the higher the decision inaccuracy. For the sake of simplicity, decision inaccuracy is assumed to be a sigmoid function of task difficulty and RT (Fig 2f). Hence, it is plausible to estimate decision inaccuracy and decision uncertainty in the mentalizing tasks merely from the external information. However, one indispensable process to distinguish mentalizing from non-social inferences or associations is the target subject’s perspective taking. For this purpose, in the type-1 mentalizing tasks (AO-DI and PS-DI), the participant should consider that the target subject has unique internal noise (*σ*_1_) during the perceptual decision-making process as described by the drift-diffusion model [25], which affects the target subject’s object-level performance (*i.e.*, decision inaccuracy, Fig 2f). In the type-2 mentalizing tasks (AO-DU and PS-DU), the participant should further consider that the target subject has unique internal noise (*σ*_2_) in mapping decision inaccuracy to decision uncertainty (Fig 2g), which renders the target subject’s unique metacognitive ability even with the same performance accuracy [26] (*i.e.*, a low variance results in a high metacognitive ability). We constructed the receiver-operating characteristic (ROC) curve by using the level of decision uncertainty as the criterion to judge the incorrectness of the choice in each trial, and measured the metacognitive ability as the area under the ROC curve (AUROC) indicating the extent to which the subjective ratings matched the actual decision outcomes [26] (Fig 2h). Taken together, the intrinsic mental state representations of decision inaccuracy and decision uncertainty on the target subject’s perspective should hierarchically exist in the type-1 and type-2 mentalizing tasks, respectively, beyond the associations with external information. However, the participant did not learn about the target subject’s (even for the past self) object-level and meta-level performance in the current study. Therefore, the representations of target subject’s intrinsic mental states, if they exist, are internally formed in the participant’s brain, and may not reflect the target subject’s actual intrinsic mental states.

### Behavioral results

The weights of task difficulty and RT (equation 1 in Methods) were equivalent (analysis of variance (ANOVA), task difficulty: F_[3,112]_ = 0.11, *P* = 0.95; RTs: F_[3,112]_ = 0.16, *P* = 0.92; Fig 2e), and were highly correlated across the mentalizing tasks (S3 Fig). Hence, the participant used external information equally to estimate the corresponding mental states across the mentalizing tasks. Notably, the estimates in the mentalizing tasks relied more on task difficulty than RTs (two-tailed paired t-test, t_27_ = 6.2; *P* = 2.5 × 10^−8^), while the estimates of decision uncertainty in the metacognition task relied more on RT than task difficulty (two-tailed paired t-test, t_27_ = 4.1; *P* = 5.9 × 10^−5^; Fig 2e), probably because the task difficulty (by the nature of the experimental design) were clearly discerned in the mentalizing tasks, but not in the metacognition task. The estimated decision inaccuracy/uncertainty in the mentalizing tasks was equivalently correlated with the originally-reported decision uncertainty by the AO/PS in the metacognition task (ANOVA, F_[3,112]_ = 0.28, *P* = 0.84, Fig 2f).

However, all these correlations disappeared after the associations with the external information of task difficulty and RT were regressed out (two-tailed t-test, *P*s > 0.30; Fig 2f). That is, the residuals in each of the mentalizing tasks did not further predict the actual decision uncertainty reported by the target subject in the metacognition task. Further, as the estimated decision uncertainty in each task could largely predict the actual decision outcome (true or false), we used the AUROC to characterize this consistency. On average, the AUROCs (mean: 0.67~0.71) were significantly above chance level (0.5) and showed no significant differences across all the tasks (ANOVA, F_[4,139]_ = 1.22, *P* = 0.31, Fig 2g). However, after the associations with the external information were regressed out, the residual AUROCs (measured by the estimate residuals) were no longer significantly different from the chance level in each of the mentalizing tasks (two-tailed t-test, *P*s > 0.20), but in the metacognition task it remained significant and as large as 0.62 (two-tailed t-test, t_27_ = 11.8; *P* = 3.6 × 10^−12^; Fig 2g). Thus, reliable estimates of decision inaccuracy/uncertainty in the mentalizing tasks were crucially dependent on the external information provided by the task difficulty and RT, which had stable associations with the inaccessible mental states. In striking contrast, the reported decision uncertainty in the metacognition task relied heavily on internal information that the task difficulty and RT could not explain. Similar to the previous argument in animals [16], it was hard to discern whether the mental state attributions in mentalizing were merely achieved through association between external information and the covert mental states. Nonetheless, the estimate residuals accounted for about half of the total variance in each of the mentalizing tasks, though much lower than the ratio of 0.74 in the metacognition task (two-tailed paired t-test, t_27_ = 3.5; *P* = 3.3 × 10^−4^; Fig 2h). Importantly, the residual variances in the type-2 mentalizing tasks were significantly larger than those in the type-1 mentalizing tasks (two-tailed t-test, AO: t_27_ = 2.5; *P* = 0.0096; PS: t_27_ = 2.1; *P* = 0.023). These extra variances in the type-2 mentalizing tasks were probably generated by the additional process of the target subject’s perspective-taking, as suggested by the theoretical analyses described above.

### Common neural representations of external information in mentalizing

As the external information of task difficulty and RT contributed equivalently to the estimates across the mentalizing tasks, we first tested the hypothesis that there are stable neural correlates of the two external social cues. Further, as utilizations of the external information were the same across the mentalizing tasks, the neural representations of each external cue might be also the same. To test these hypotheses, we regressed the trial-by-trial fMRI activity during the judgment phase with the order of the task difficulty and RT across the whole brain in each task (see Methods). Across all mentalizing tasks, fMRI activity in the primary visual cortex (V1) was negatively correlated with task difficulty (conjunction analysis, *z* > 2.6, *P* < 0.05 after cluster-level family-wise error (FWE) correction; Fig 3a), decreasing as the number of moving dots was reduced (increasing task difficulty). On the other hand, fMRI activity in the right IPL was positively correlated with task difficulty (conjunction analysis, *z* > 2.6, *P* < 0.05 after cluster-level FWE correction; Fig 3a). In contrast, the fMRI activity in a wide range of brain regions was positively correlated with RT (conjunction analysis, *z*> 2.6, *P* < 0.05 after cluster-level FWE correction; Fig 3b). Among these brain regions, the right IPL region also overlapped with the regions associated with task difficulty: the same right IPL region responded to both task difficulty and RT in the mentalizing tasks (Fig 3c and 3d). Thus, integration of the two pieces of external information in the right IPL partially predicted decision inaccuracy/uncertainty.

**Fig 3.**
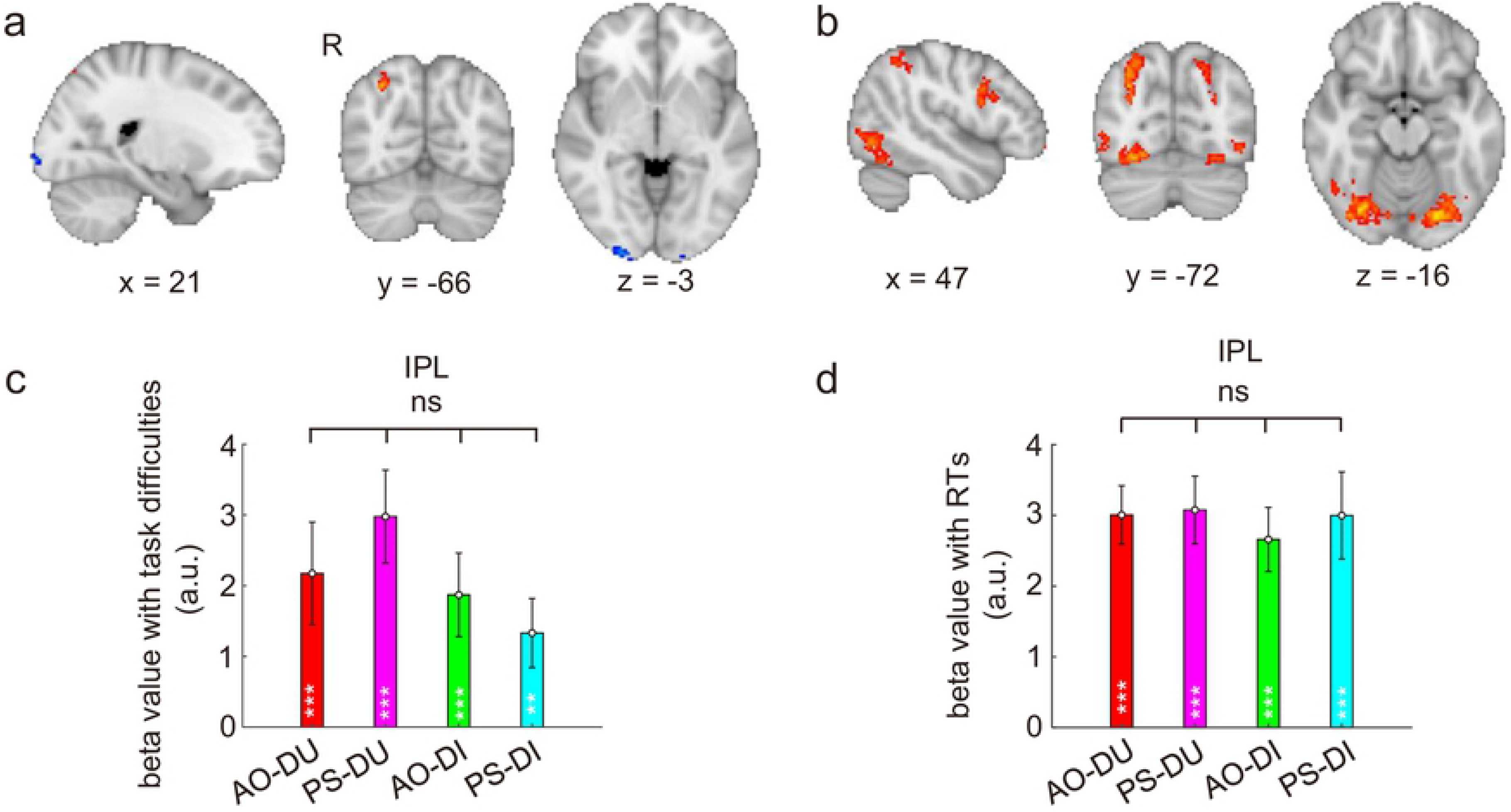
Neural representations of external information common across the mentalizing tasks. (**a**) The activation maps for the activities significantly correlated with task difficulty in a conjunction analysis across the four mentalizing tasks (AO-DU, PS-DU, AO-DI, PS-DI) (*z* > 2.6, *P* < 0.05 after cluster-level FWE correction). (**b**) The activation maps for the activities significantly correlated with RT in a conjunction analysis across the four mentalizing tasks (*z* > 2.6, *P* < 0.05 after cluster-level FWE correction). (**c**) The parametric regression beta values of task difficulty in each of the mentalizing tasks in the right IPL ROI defined by the conjunction of (**a**) and (**b**). (**d**) The beta values of RT in each of the mentalizing tasks in the right IPL ROI defined by the conjunction analysis of (**a**) and (**b**). The error bars represent SEM across participants. *ns*, not significant; \**P* < 0.05; \*\**P* < 0.01; \*\*\**P*< 0.001.

### Distinct neural representations of estimate residuals in mentalizing

According to our theoretical analyses as described above, additional unique processes should be involved in each of the mentalizing tasks besides the associations with external information. To explore the underlying neural correlates, we regressed the estimate residuals with trial-by-trial fMRI activities during the judgment phase across the whole-brain voxels in each task. The modulation effects were prevalent during the judgment phase by comparison to the alternative GLM accounting for the modulation effects during the decision-making phase (S2 Fig).

In the type-2 mentalizing task, where the participant estimated the AO’s decision uncertainty (AO-DU), the estimated residuals were significantly correlated with the fMRI activities in the dmPFC, dorsally and anteriorly neighboring, but separate from, the dACC region representing the current-self decision uncertainty in metacognition (see below), as well as in the left TPJ and the left inferior frontal junction (IFJ) (*z* > 3.1, *P* < 0.05 after cluster-level FWE correction, Fig 4a; see also S1 Table). In the type-1 mentalizing task, where the participant estimated the AO’s decision inaccuracy (AO-DI), the estimated residuals were also correlated with the fMRI activities in the left TPJ and the left IFJ (*z* > 3.1, *P* < 0.05 after cluster-level FWE correction, Fig 4b and 4e; see also S1 Table), but not in the dmPFC (t_27_ = 1.1, *P* = 0.12; Fig 4e), suggesting that dmPFC was selectively involved in type-2 mentalizing. To further test the reliability of the dmPFC selectivity in type-2 mentalizing, we repeated the same GLM analysis on the dmPFC and TPJ regions independently defined by meta-analytical maps from the NeuroSynth database [27], as well as the conjunction regions between the meta-analytical regions and those in the current study. In both analyses, we obtained results consistently supporting that dmPFC but not TPJ showed activity selective to type-2 mentalizing (S4 Fig).

**Fig 4.**
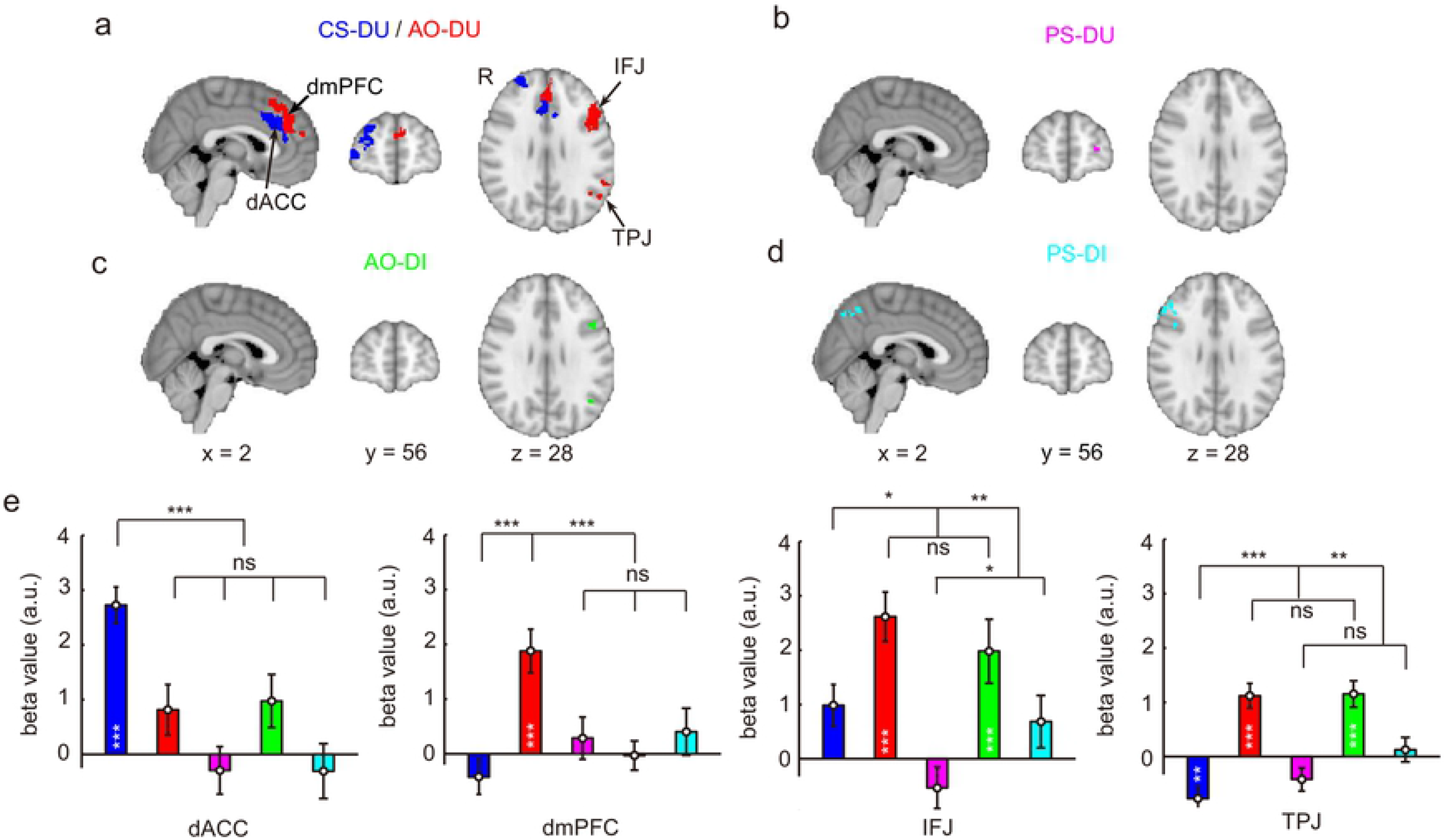
Distinct neural representations of estimate residuals in metacognition and mentalizing. (**a**) Neural correlates of the estimated residuals of decision uncertainty in the metacognition task (dACC and lFPC, blue) and the estimated residuals of decision uncertainty in the AO-DU (type-2) mentalizing task (dmPFC and left-IFJ and left-TPJ, red). (**b**) Neural correlates of the estimate residuals of decision uncertainty in the PS-DU (type-2) mentalizing task (left lFPC, magenta). (**c**) Neural correlates of the estimate residuals of decision inaccuracy in the AO-DI (type-1) mentalizing task (left-IFJ and left-TPJ, green). (**d**) Neural correlates of the estimate residuals of decision inaccuracy in the PS-DI (type-1) mentalizing task (right-IFJ and precuneus, cyan). (**e**) The comparisons of the parametric regression beta values with the estimate residuals across tasks in the ROIs of dACC, dmPFC, left-IFJ and left-TPJ (see also S4 Fig). The error bars represent SEM across participants. *ns*, not significant; \**P* < 0.05; \*\**P* < 0.01; \*\*\**P*< 0.001.

Instead, the estimate residuals of the PS decision uncertainty in the type-2 mentalizing task were selectively correlated with the fMRI activities in the right lFPC (*z* > 3.1, *P* < 0.05 after cluster-level FWE correction, Fig. 4c and S5f Fig), whereas the estimated residuals of the PS decision inaccuracy were correlated with the fMRI activities in the precuneus and the right IFJ in the type-1 mentalizing task (*z* > 3.1, *P* < 0.05 after cluster-level FWE correction, Fig. 4d and S5e Fig). Critically, both the right lFPC and precuneus regions are also shared with the metacognition-associated areas reported in prior studies [18, 19].

Consisting with our theoretical account, selective neural representations of estimate residuals in the mentalizing tasks showed that these residuals were partially but reliably correlated with diverse brain activities during mentalizing in different contexts. Notably, the right IPL that encoded task difficulty and RT did not represent the estimate residuals in the mentalizing tasks (S5a Fig). The dissociation of neural representations of external information from the estimate residuals confirms that the intrinsic mental state representations do coexist in mentalizing to attribute mental states to the target subjects other than the current self.

### Neural representations of estimate residuals in metacognition

In the metacognition task (CS-DU), the estimated residuals were significantly correlated with the fMRI activities in the dACC and the lFPC (*z* > 3.1, *P* < 0.05 after cluster-level FWE correction, Fig 4a; see also S1 Table), as repeatedly observed in previous studies [18, 19, 28]. Although the lFPC region was shared with type-2 mentalizing (PS-DU), the dACC region selectively represented the estimate residuals in the metacognition task but not in the mentalizing tasks. Notably, the components of decision uncertainty associated with task difficulty and RT in the metacognition task were also represented in the dACC (S5b Fig). Thus, the dACC uniformly represented the components of internally-generated decision uncertainty in metacognition.

### Common neural representations of estimate uncertainty in mentalizing

In the mentalizing tasks, the use of associations with external cues could not provide sufficient information for the trial-by-trial estimation of AO/PS decision inaccuracy/uncertainty. Further, the intrinsic mental state attributions also did not rely on the target subject’s attributes that were unknown for the participant in the current study. Thereby, the estimating processes often brought out estimate uncertainty [29, 30]. The estimates were usually more uncertain at the middle levels: longer RTs in reporting ratings at the middle levels than at the lowest and highest levels (inverted U-shape, Fig 5a). We divided all the trials equally into eight bins according to the quantity of external information, calculated by a sigmoid function of the task difficulty and RT (equation 1 in Methods). RTs were also longer at the middle bins than at the lower and higher bins (inverted U-shape, Fig 5b). The estimate uncertainty was the second-order variable of the estimates, similar to decision uncertainty about the decisions in metacognition. We then operationally defined the trial-by-trial estimate uncertainty as - |uncertainty-mean (uncertainty)|. We regressed the estimate uncertainty with the trial-by-trial fMRI activities after the associations with decision inaccuracy/uncertainty were regressed out in each task. The neural correlates of estimate uncertainty in the mentalizing tasks were similar to those of decision uncertainty in metacognition (S6 Fig). Across the mentalizing tasks, the fMRI activity in the dACC was significantly and commonly correlated with estimate uncertainty (Fig 5c). Critically, the dACC region associated with estimate uncertainty across the mentalizing tasks was largely overlapping with the dACC region associated with decision uncertainty in the metacognition task (Fig 5d).

**Fig 5.**
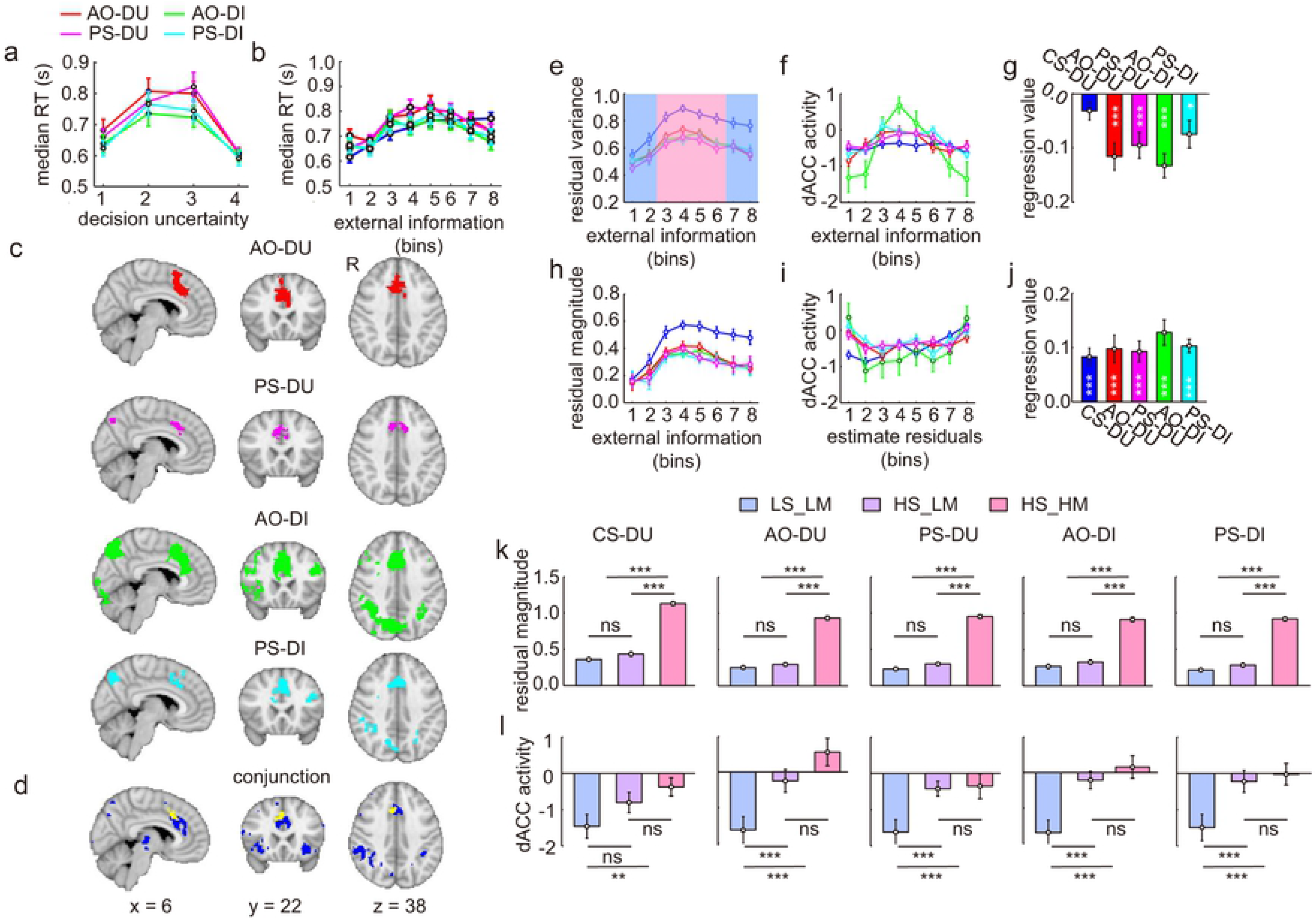
The dACC activities represented estimate uncertainty across the mentalizing tasks. (**a**) The RT for reporting the estimates was longer at the middle levels of decision inaccuracy/uncertainty; (**b**) The RT for reporting the estimates was longer at the middle levels of external information calculated by a sigmoid function of task difficulty and RT, than at the lower and higher levels. (**c**) The neural activities were significantly correlated with the estimate uncertainty (−1*|uncertainty –mean(uncertainty)|) in each of the mentalizing tasks. (**d**) The dACC was commonly involved in monitoring estimate uncertainty (conjunction analysis, yellow), overlapping with monitoring the current-self decision uncertainty in the metacognition task (blue, see also Fig 4a). (**e**) The residual variances were larger at the middle levels of external information. (**f**) The dACC activity was larger at the middle levels of external information. (**g**) The parametric regression beta values of the dACC activity with absolute values of external information. (**h**) The residual magnitudes were larger at the middle levels of external information. (**i**) The dACC activity was larger when the residual magnitudes were larger. (**j**) The regression beta values of the dACC activity with the residual magnitudes. (**k**) The trials were divided into three sub-groups with the same residual variances but different residual magnitudes (HV_LM: high_variance-low_magnitude vs. HV-HM: high_variance-high_magnitude) and the same residual magnitudes but different residual variances (HV-LM vs. LV_LM: low_variance-low_magnitude). (**l**) The dACC activity was significantly different between different residual variances, but not between different residual magnitudes. The error bars represent SEM across the participants. *ns*, not significant; \**P* < 0.05; \*\**P* < 0.01; \*\*\**P*< 0.001.

According to estimation theory [29, 30], the variance of the estimate residuals should be larger in the central range of the external information (Fig 2f). Exactly consistent with this prediction, the variance of the estimate residuals in each bin of the external information was a negative parabolic function in each task (inverted U-shape, Fig 5e). Accordingly, the mean dACC activity in each bin of the external information was also a negative parabolic function in each of the mentalizing tasks (inverted U-shape, *P*s < 0.05; Fig 5f and 5g), but not in the metacognition task (two-tailed t test, t_27_ = −1.9; *P* = 0.07; Fig 5g). Altogether, these results suggest that the dACC also plays an important role in monitoring the mentalizing process. Notably, this is complementary with prior findings that activity in the ventromedial prefrontal cortex (vmPFC) and the posterior cingulate cortex (PCC) are responsive to estimate confidence [29].

### dACC was involved in monitoring the estimate variance, rather than in generating the estimates

Larger residual variances in the central bins might also reflect larger residual magnitudes. To examine this potential confound, we calculated the mean residual magnitude averaged in each bin of the external information also sorted by the quantities calculated by the sigmoid function of task difficulty and RT. The residual magnitudes had a similar relationship with the bins as the residual variances did (Fig 5h). Accordingly, the dACC activity in each bin of the estimate residuals according to the residual values (the mean was zero) had a positive parabolic function in each of the tasks, including the metacognition task (U-shape, *P*s < 0.001; Fig 5i and 5j). That is, the dACC activity increased as the residual magnitude increased.

Since the dACC has also been suggested to play a critical role in control of flexible adaptation [31–33], it remains possible that this region played a control role in generating the estimate residuals, rather than in monitoring the estimate variances. This possibility was also evident by the observation that larger residual magnitude was associated with greater dACC activity (Fig 5i). To clearly identify the functional role of the dACC in mentalizing, we first median-split the trials on residual variances (Fig 5e, low variances: blue zone, high variances: red zone). Accordingly, the residual magnitudes should be also low in the former group (LV_LM: low variances and low magnitudes), whereas the residual magnitudes were further segregated into low (HV_LM: high variances and low magnitudes) and high (HV_HM: high variances and high magnitudes) sub-groups in the latter group by another median-split (Fig 5k). By virtue of these two divisions, although the residual magnitudes were not different between the LV_LM and HV_LM conditions, their variances were significantly different (*P*s < 6.7 × 10^−14^; Fig 5k). In contrast, although the residual magnitudes were significantly different between the HV_LM and HV_HM conditions, their variances were not different (*P*s > 0.18; Fig 5k). Hence, we tested whether the dACC activities were selectively responsive to the residual variances or the residual magnitudes by comparing the dACC activities among the three sub-groups. If the dACC activities were sensitive to the residual variances, but not residual magnitudes, then it should be significantly different between the LV_LM and HV_LM conditions, but not between the HV_LM and HV_HM conditions. Otherwise, the dACC activity pattern should be reverse among the three sub-groups. The results demonstrated that the dACC activity pattern concurred well with the former prediction across the mentalizing tasks (Fig 5l), supporting that the dACC was involved in monitoring the residual variances (estimate uncertainty), rather than in generating these residuals. That is, metacognition monitors mentalizing (Fig 6a). Importantly, the same format of mental state representations of decision uncertainty in metacognition and estimate uncertainty in mentalizing were registered in the dACC. On the contrary, mentalizing was not subject to metacognition and was not associated with the dACC activities. Instead, different forms of mentalizing had diverse formats of mental state representations of inferred decision uncertainty in the human brain (Fig 6b).

**Fig 6.**
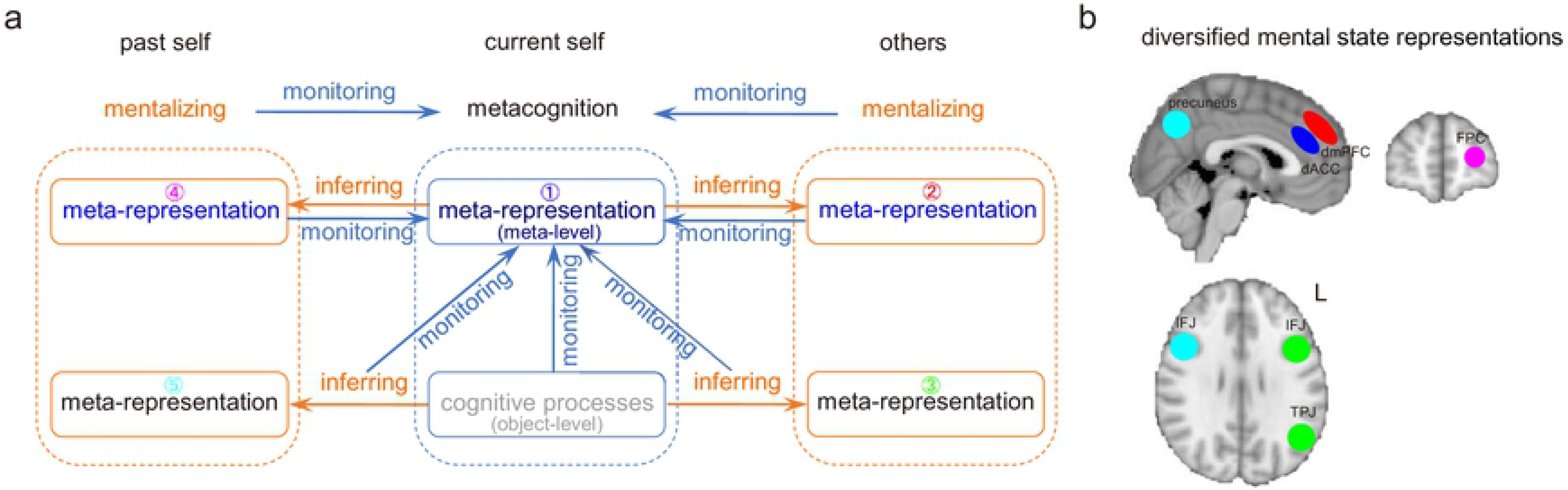
The neural representations of decision uncertainty in mentalizing and metacognition. (**a**) The distinctions between metacognition and mentalizing rely on whether the monitored information is accessible, rather than whether the target subject to be attributed is the self or others. Hence, attributing mental states to the past self is also mentalizing. Further, although meta-representations in metacognition and mentalizing are nearly separate, metacognition also accompanies and monitors all the mentalizing processes. (**b**) Distinct neural representations of mental states in metacognition and mentalizing. The left IFJ and left TPJ represent the other’s decision inaccuracy (mental construct of the object-level performance) and the dmPFC represents the other’s decision uncertainty (mental construct of the metacognitive performance). In contrast, the precuneus and the right IFJ represent past-self decision inaccuracy, and the lFPC represents past-self decision uncertainty. In contrast, the dACC commonly represents the current-self decision uncertainty in metacognition and estimate uncertainty in mentalizing.

## Discussion

The relationship between metacognition and mentalizing is still a matter of debate [34]. In the current study, we adapted an experimental paradigm from metacognition to mentalizing. We could thus parametrically distinguish the neural representations of the mental states of decision uncertainty in both metacognition and mentalizing. Using fMRI to characterize the neural signals correlated with the corresponding mental states, we identified for the first time the different mental state representations underlying the attribution of decision inaccuracy/uncertainty to different targets (the current self, the past self, and others), independent of external information. These separate mental state representations of decision uncertainty clearly distinguish mentalizing from metacognition.

Metacognition in the current study involved in monitoring internally-generated decision uncertainty. Behaviorally, even after the perceptual information of task difficulty and RT was regressed out, the residuals stably predicted the decision outcomes, as these mental states were internally accessible [1–3]. Neurally, the dACC represented these components of decision uncertainty. In contrast, mentalizing in the current study involved monitoring inferred decision uncertainty. The participant could only use external information to infer the inaccessible mental states. Unsurprisingly, after the perceptual information was regressed out, the estimate residuals did not further predict the target subject’s actual mental states of decision uncertainty. Nonetheless, stable neural representations of these estimate residuals were reliably observed in the mentalizing tasks. Critically, these neural representations of estimate residuals in attributing different levels (object-level and meta-level) of mental states to different targets (AO and PS) were distinct. Our empirical results thus confirm our theoretical account: mentalizing recruits additional mental processes beyond associations with external information. Critically, these intrinsic mental state representations made essential distinctions of mentalizing in different contexts.

The type-1 mentalizing task in monitoring an anonymous other’s decision inaccuracy (AO-DI) was similar to false-belief tasks (*e.g.*, the “Sally-Anne” task) [4, 35, 36]. Common to both tasks, the participant needs to judge the other’s performance, that is, the choice probability of the incorrect option, or decision inaccuracy (Sally should always choose the incorrect option in the “Sally-Anne” task), while the participant actually knows the truth. To attribute object-level states to the other, the participant parsimoniously makes a counterfactual inference on the basis of stable associations with external cues but does not necessarily construct a model for the other’s mental world. According to the theoretical account, to take the other’s perspective, the participant merely builds up a simple model describing the other’s task performance (type-1) with common knowledge that each person should have a unique capability in task performance (*e.g.*, internal noise *σ*_1_) even with the same stimulus information. Our results imply that activity in the left TPJ and the left IFJ might be associated with this mental process. This notion is consistent with prior evidence that the TPJ activation is saliently observed in false-belief tasks [21, 35, 36], while anatomical and virtual lesions in the TPJ region selectively cause serious deficits in perspective taking [37, 38].

The AO-DU type-2 mentalizing task was identical to the AO-DI type-1 mentalizing task except that the participant estimated the AO’s meta-level mental state of decision uncertainty about the AO’s own performance. Hence, beyond the mental processes involved in the type-1 mentalizing task, the participant needed a model for the AO’s metacognitive ability. Our results imply that this mental model might be built in the dmPFC, which was selectively activated in the type-2 mentalizing task, but not in the type-1 mentalizing task. Although both the dmPFC and the TPJ have often been shown to be involved in mentalizing [7, 21, 39], our findings specify a functional distinction. The dmPFC was specifically involved in constructing the mental model of the other’s meta-level mental states [40, 41]. In striking contrast, the TPJ might be generally involved in representing the other’s object-level mental states. Crucially, when the identical task sequence was used but the target subject was changed from an anonymous other to the past self, the neural loci associated with the estimate residuals of decision inaccuracy/uncertainty were altered. Thereby, these specific other-oriented neural representations of decision uncertainty in the dmPFC and TPJ regions support the theory of mind (ToM) in accounting for mentalizing [7, 21, 39].

When the participant estimated the past-self decision inaccuracy, activity in the precuneus and the IFJ was selectively associated with the estimate residuals. On the other hand, when the participant estimated the past-self decision uncertainty, the activities in the lFPC and the IFJ were selectively associated with the estimate residuals. The lFPC and precuneus regions are also both associated with metacognitive processes in prior studies [18, 19]. Hence, these results suggest that mentalizing for the past-self mental states of decision uncertainty recruited neural loci shared with metacognition. In other words, these self-oriented neural representations of decision uncertainty in the lFPC and the precuneus supported the simulation theory (ST) in accounting for mentalizing [42]. However, the dACC region, the crucial brain region for monitoring the current-self mental states of decision uncertainty [18, 28], was not activated in the past-self-oriented mentalizing tasks (S5b Fig).

When the participant monitored the AO/PS’s task performance in the type-1 mentalizing task, similar to monitoring the current-self task performance in the metacognition task, the reported AO/PS’s decision inaccuracy was the subjective beliefs of the participant, rather than the target subject. However, as described above, the neural representations of these mental states in the type-1 mentalizing tasks were entirely different from those in the metacognition task. To this end, the critical distinction between metacognition and mentalizing should depend on the accessibility of sources to be monitored, rather than the agents to whom the mental states are tied or the target subjects to whom the mental states are attributed. Altogether, our results illustrate that the human brain diversifies separate neural systems to represent the different mental states of decision uncertainty in monitoring the current self, the past self, and others in performing the same perceptual decision-making task.

Importantly, our current neuroimaging results in particular clearly illustrate the relationship between metacognition and mentalizing, a longstanding puzzle in psychology and philosophy. First, the mental state representations in metacognition and mentalizing are dissociated. Metacognition and mentalizing are two independent processes with different meta-representations. Second, metacognition accompanies and monitors mentalizing, but mentalizing is not necessarily dependent on metacognition. The dACC commonly monitors estimate uncertainty across the different mentalizing tasks and decision uncertainty in metacognition. In essence, mentalizing is a perception-based social inference process. Hence, metacognition as a domain-general process should also monitor this type of high-level cognitive processes [18]. Third, even though the participant did not explicitly report estimate uncertainty during mentalizing, the dACC could implicitly or automatically capture estimate uncertainty [21].

Mentalizing is a crucial social cognitive function for human behaviors. During interpersonal interactions, the primary motivation of mentalizing is to predict and influence others’ beliefs, desires and intentions, as well as their actions [41]. It is plausibly an effective strategy to manipulate influences on others when they are uncertain, rather than when they are highly confident, since the odds of success in changing others’ minds should be higher in the former case. Even for preverbal infants, when they feel uncertain, they are willing to seek caregiver’s helps [43]. Therefore, across the mentalizing tasks, the target subject’s decision inaccuracy/uncertainty, rather than the decision accuracy/confidence, were predominantly positively correlated with the brain activities, which might be used to guide appropriate social control [44, 45]. Hence, the dACC involved in monitoring the current-self decision uncertainty in metacognition and the dmPFC involved in monitoring others’ decision uncertainty in mentalizing drive cognitive control and social control, respectively. The dACC region here is anatomically in the sulcus of ACC (ACCs), while the dmPFC region is dorsally neighboring dACC [46]. Prior studies have also shown that a region in the gyrus of ACC (ACCg) or the perigenual ACC (pgACC), ventrally neighboring the dACC, is also involved in monitoring and predicting the behaviors of the others [47], but specifically in tracking the motivational values of the others [48].

However, estimates of others’ mental states and even those of the past-self mental states were not predictable for the target’s intrinsic mental states and performance (after the associations with external information were regressed out). These findings thus implicate that volatile momentary mental states are quite difficult to predict, probably due to the fact that the AO’s or PS attributes in both object-level and meta-level performance were unknown to the participant in the current study. One potential approach to improve the predictability of mentalizing could be through social learning from interpersonal interactions [45], to construct more reliable mental models for a specific target’s mental world from more subtle social information and social experience [49–51]. Meanwhile, metacognition might facilitate social learning processes by monitoring and controlling mentalizing. Our findings of multiple mental state representations in mentalizing for different targets under different levels of sources provide new insight on neural computations of the internal mental models in different mentalizing processes. Eventually, these may improve future diagnostic approaches for social deficits in ASD and schizophrenia.

## Methods

### Participants

We recruited twenty-eight healthy right-handed participants (22 females, age: 23.5 ± 1.5 years old) to take part in all five tasks across three fMRI experiments. Informed consent was obtained from each participant in accordance with a protocol approved by Beijing Normal University Research Ethics Committee (ICBIR_A_0091_004).

### Metacognition (random dot motion: RDM) task

In an aperture with a radius of three degrees of visual angle, three hundred white dots (radius: 0.08 degrees, density: 2.0%) were displayed on a black background. The dots moved in different directions at a speed of 8.0 degrees/second. The movement of each dot lasted three frames. A subset of dots moved coherently in the same direction (left, up, right, or down), while the other dots moved in different random directions. The participant was required to discriminate the net motion direction and rate her uncertainty about the decision by pressing a corresponding button. The task difficulty was determined by the percentage of coherently moving dots, with four levels of task difficulty randomly mixed in the task. The easiest and hardest levels were fixed at a coherence of 1% and 30%, respectively, while the two middle levels were set for each participant to achieve accuracy of 50% and 80%, determined by a staircase procedure in a practice session conducted several days prior to the fMRI experiments [18, 28].

### Mentalizing tasks

A pair of participants who had similar stimulus coherences at the 50% and 80% accuracy level in the practice session jointly took part in the task (the stimulus coherences at the two task difficulty levels used in the mentalizing tasks were the means of their stimulus coherences, respectively). One (the target subject) performed the RDM task outside the scanner, and another (the participant) observed the target’s performance from inside the scanner. The two computers separately presented the stimuli but were connected and synchronized by the local network following the TCP/IP protocol. Hence, they simultaneously performed their own tasks with the same sequence (see below) but in response to different stimuli and task requirements. To avoid eliciting her own decision uncertainty, the stimulus presented to the participant inside the scanner was noiseless: the randomly-moving dots remained stationary and only the coherently moving dots moved. While the target subject was responding, the participant saw a progress bar representing the elapsed time. However, the participant could not see the target’s choice and the reported decision uncertainty. The participant then reported the estimate of the target subject’s decision inaccuracy/uncertainty. In the PS-DU and PS-DI mentalizing tasks, the target subject was replaced by the past self. That is, the participant observed RDM task performance previously done by herself.

### Task sequence

The sequences of the metacognition and mentalizing tasks were almost identical. In the metacognition task, each trial started with a green cross cue to indicate that the task stimulus would be presented 1 s later. The stimulus was then presented for 2 s, and four options of the moving directions were presented. The participant made a choice within 2 s. After a choice was made, four ratings from 1 (most uncertain) to 4 (most certain) were presented, and the participant reported the rating by pressing the corresponding button within 2 s. The inter-trial interval (ITI) was jittered uniformly between 2 s and 6 s. On average, each trial lasted for 9 s. During the mentalizing tasks, although the sequence was identical to the metacognition task performed outside the scanner, the participant watched the noiseless stimulus presentation and the progress bar indicating elapsed time for the target’s response, and then reported the estimate rating of the target’s decision inaccuracy/uncertainty. Each task was conducted for two consecutive runs of sixty trials. The task order of the three experiments was random and counterbalanced across the participants.

### Behavioral analyses

The reported ratings of decision inaccuracy/uncertainty in each of the metacognition and mentalizing tasks were associated with the task difficulty and RT. We used a sigmoid function to characterize the relationship as follows,

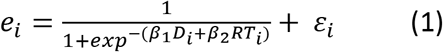

where *e*_*i*_ represents the estimate of decision inaccuracy/uncertainty at the trial *i*, *D*_*i*_ and *RT*_*i*_ represents the task difficulty and the response time at the trial *i*, respectively, and *ɛ*_*i*_ represents the estimated residual. Task difficulties and RTs were separately normalized within each participant. Further, for the mentalizing tasks, we also considered an autoregressive (AR) model to account for the associations of the estimates between the contiguous trials. However, the autocorrelation of the estimates between the contiguous trials in each mentalizing task was not significant, indicating that the estimates were not dependent on the history, but only on the current trial.

### fMRI parameters

All fMRI experiments were conducted using a 3-T Siemens Trio MRI system with a 12-channel head coil (Siemens, Germany). Functional images were acquired with a single-shot gradient echo T_2_^*^ echo-planar imaging (EPI) sequence with volume repetition time of 2 s, echo time of 30 ms, slice thickness of 3.0 mm and in-plane resolution of 3.0 × 3.0 mm^2^ (field of view: 192 × 192 mm^2^; flip angle: 90 degrees). Thirty-eight axial slices were taken with interleaved acquisition, parallel to the anterior commissure-posterior commissure line.

### fMRI analyses

fMRI analyses were conducted using FMRIB’s Software Library (FSL) [52]. To correct for rigid head motion, all EPI images were realigned to the first volume of the first scan. Data sets in which translation motion was larger than 2.0 mm or rotation motion was larger than 1.0 degree were discarded. No data were discarded in these analyses. Brain matter was separated from non-brain using a mesh deformation approach and used to transform the EPI images to individual high-resolution structural images, and then to Montreal Neurological Institute (MNI) space by using affine registration with 12 degrees of freedom and resampling the data with a resolution of 2 × 2 × 2 mm^3^. Spatial smoothing with a 4-mm Gaussian kernel (full width at half-maximum) and high-pass temporal filtering with a cutoff of 0.005 Hz were applied to all fMRI data.

We used generalized linear modeling (GLM) to analyze the fMRI data. For the first-level analyses, two events were modeled in each trial. The first event (the stimulation phase) represented the stimulus presentation, time-locked to the onset of the stimulus presentation, with a duration of the presentation time (2 s). The second event (the judgment phase) represented estimating decision inaccuracy/uncertainty, time-locked to the onset of the rating, with the duration of the reaction time. The parametric modulation effects of task difficulty and RT, the estimate residual, and the estimate uncertainty (−1*|uncertainty – mean(uncertainty)|) were simultaneously added in the latter event (the judgment phase). The estimate residual was the residual after the components associated with the task difficulty and RT were regressed out. All the regressors were convolved with the canonical hemodynamic response function with double-gamma kennels. The results when the parametric modulation effects were added in the stimulation phase are presented in S2 Fig.

For the group-level analyses, we used FMRIB’s local analysis of mixed effects (FLAME), which models both ‘fixed effects’ of within-participant variance and ‘random effects’ of between-participant variance using Gaussian random-field theory in each task. Statistical parametric maps were generated with a threshold of *z* > 3.1, *P* < 0.05 after cluster-level family-wise error (FWE) correction for multiple comparisons for each contrast, unless mentioned otherwise.

### Conjunction analyses

We conducted conjunction analyses to test the common neural correlates of external social cues (task difficulty and RT) across the four mentalizing tasks (AO-DI, PS-DI, AO-DU, PS-DU), as well as the common neural correlates of the estimate uncertainty across the four mentalizing tasks. We identified significant regions in which there was evidence of effects in all contrasts of conditions using the FSL script *easythresh*. Statistical parametric maps were generated by the threshold with *z* > 2.6, *P* < 0.05 after cluster-level FWE correction for each contrast.

### Region-of-interest (ROI) analyses

To circumvent circular inference, we randomly analyzed the data from three fourths of the participants to define each of the ROIs with a significance of z > 2.6, *P* < 0.05 after cluster-level FWE correction, and then used the data from the held-out participants to obtain the parametric regression beta value in each ROI. We repeated this analysis one hundred times and averaged the parametric regression beta values for each ROI. For the dmPFC and TPJ ROIs, we further assessed meta-analytically-derived ROIs associated with from Neurosynth (search term: ‘mentalizing’) [27], as well as the overlapping regions between our results and the Neurosynth ROIs. For the dACC region in metacognition, we used the overlap between the RDM task and the mentalizing tasks obtained by conjunction analysis.

To identify whether the dACC activities in mentalizing tasks were associated with the residual variances (*i.e.*, monitoring estimate uncertainty) or the residual magnitudes (*i.e.*, generating the residuals), we sequentially divided the trials in each of the metacognition and mentalizing tasks into three sub-groups according to the residual variances and the residual magnitudes. First, we split the trials into two sub-groups with low and high residual variances according to whether the residual variance was above the median, and further split the trials with high residual variances into two sub-groups with low and high residual magnitudes similarly according to whether the residual magnitude was above the median (Fig 5j). We then calculated and compared the dACC activity between the sub-groups (Fig 5k).

### Simulations and parameter recovery

Because the stimulus presentation and uncertainty rating events were temporally close to each other, the GLM estimates for the two events should be highly collinear. To examine whether the GLM analyses could separate the underlying neural activity specifically associated with each of the two events, we generated fMRI data using the different parameters of the GLM as follows and validated the parameter recovery [27].

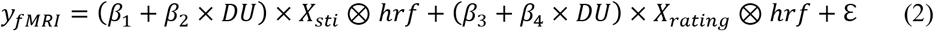

where X_sti_ and X_rating_ represent the design matrix for the stimulation phase and the judgment phase, respectively. *β*_1_ and *β*_3_ are the mean activities of the two events, *β*_2_ and *β*_4_ are the estimate uncertainty modulation effects on the two events, respectively. *hrf* is the canonical hemodynamic response function with two-gamma kennels, ε is an additional Gaussian noise. The values of *β*_1_ and *β*_3_, as well as *β*_2_ or *β*_4_ were independently and randomly drawn from a uniform distribution in the range of [0.2,0.8], while the alternative *β*_2_ or *β*_4_ was set to zero. That is, the modulation effect appeared only in either of the two phases, but not simultaneously in both phases. The signal-to-noise (SNR) of the fMRI data were set uniformly in the range of [0.01, 1] (the event-evoked fMRI signals are usually in the range of 0.1~10 of the noise). We then used the GLM with the mean activity and parametric modulation in both phases to reconstruct the mean activity and the parametric modulation effects in the two events from the generated fMRI data with the sampling rate of 0.5 Hz (TR = 2 s). For each set of the parameters, 1,000 times of procedures were repeated and the estimated values were averaged at each SNR level (S2 Fig).

## Acknowledgments

This research was funded by the Key Program for International S&T Cooperation Projects of China (MOST, 2016YFE0129100, X.W.), the National Natural Science Foundation of China (No. 31471068, X.W.), the Fundamental Research Funds for the Central Universities (2017EYT33, X.W.).

## Data availability

All the data and codes used in the current study are available in https://github.com/ShaohanJiang/UncertaintyinMetacogMetalizing.git.

## Supporting Information

**S1 Fig. Performance accuracy and response time (RT) changed with task difficulty in the RDM task.** The four task difficulty levels for each individual participant were set at 1% coherence, 50% accuracy (actually 55% inside the scanner), 80% accuracy, and 30% coherence, respectively (chance accuracy level: 25%). The error bars represent SEM across participants.

**S2 Fig. Reliable separation of neural activity during the stimulation phase and the judgment phase (GLM analyses).** The fMRI signals were simulated by the GLM as shown, taking into account the separate contributions from the two phases, where DU represents the modulation effect of decision uncertainty, Ɛ represents the Gaussian noise, and hrf represents the canonical hemodynamic response function. The fMRI signals were obtained by the GLM with different mean values (0.2~0.8) simultaneously at both phases and the decision uncertainty modulation component (0.2~0.8) was added into the stimulation phase or the judgment phase. However, The recovered parameters were fitted by the GLM with the parametric modulation effects in both phases. For each set of parameters, procedures were repeated 1,000 times and the estimated values were averaged at each SNR level. (**a**) The correlation between the components of generated fMRI time series in the stimulation phase and in the judgment phase in each participant (the event sequence was identical to the AO-DU task). (**b**) The ratio of the recovered mean activity (*β*_1_ and *β*_3_) to the original mean activity in each corresponding phase under the different SNR levels. (**c**) The recovered parametric modulation effects (*β*_2_ in the stimulation phase and *β*_4_ in the judgment phase) under the different SNR levels. The original mean activities were in both phases but the parametric modulation effect was in the stimulation phase in (**b**) and (**c**). (**d**) The ratio of the recovered mean activity (*β*_1_ and *β*_3_) to the original mean activity in each corresponding phase under the different SNR levels. (**e**) The recovered parametric modulation effects (*β*_2_ in the stimulation phase and *β*_4_ in the judgment phase) under the different SNR levels. The original mean activities were in both phases but the parametric modulation effect was in the judgement phase in (**d**) and (**e**). (**f**)-(**i**) The parametric regression beta values in the GLM for the modulation effects at the stimulation phase for the empirical data. Except for the dACC in the CS-DU task, the modulation effects were not significant in the other tasks or in the other ROIs. The error bars represent SEM across participants. \*\*\**P* < 0.001.

**S3 Fig. Relationships between the weights of task difficulty and RT in accounting for the estimates across tasks.** The weights of task difficulty and RT were highly correlated and equivalent (falling along the diagonal) across the mentalizing tasks. However, the weights of the task difficulty were larger in the mentalizing tasks than in the metacognition task, but the weights of the RTs were reversed.

**S4 Fig. Neural parametric regression beta values with the estimate residuals in each task.** (**a**) The meta-analytically derived ROIs associated with ‘mentalizing’ from Neurosynth; (**b**) The overlapped regions between our results and the meta-analytically-derived ROIs. The convention of the colors is the same as in Fig 4. The error bars represent SEM across participants. *ns* not significant; \**P* < 0.05; \*\**P* < 0.01; \*\*\**P* < 0.001.

**S5 Fig. The parametric beta values of fMRI activity regressed with task difficulty, RT, and estimate residuals in each task in the ROIs.** (**a**) IPL; (**b**) dACC; (**c**) TPJ (**d**) dmPFC; (**e**) precuneus; (**f**) FPC. The convention of the colors is the same as in Fig 4. The error bars represent SEM across participants. \**P* < 0.05; \*\**P* < 0.01; \*\*\**P*< 0.001.

**S6 Fig. Activation maps showing significantly correlated with the estimate uncertainty in each task.** (**a**) CS-DU; (**b**) AO-DU; (**c**) PS-DU; (**d**) AO-DI; (**e**) PS-DI. *z* > 3.1, *P* < 0.05, after cluster-level FWE correction.

**S1 Table. Brain activations summary.**

## Notes

**Competing Interest Statement:** There were no conflict interests.

### Competing Interest Statement

The authors have declared no competing interest.

